# Molecular characterization of breast cancer cell response to metabolic drugs

**DOI:** 10.1101/185082

**Authors:** Lucía Trilla-Fuertes, Angelo Gámez-Pozo, Jorge M Arevalillo, Mariana Díaz-Almirón, Guillermo Prado-Vázquez, Andrea Zapater-Moros, Hilario Navarro, Rosa Aras-López, Irene Dapía, Rocío López-Vacas, Paolo Nanni, Sara Llorente-Armijo, Pedro Arias, Alberto M. Borobia, Paloma Maín, Jaime Feliú, Enrique Espinosa, Juan Ángel Fresno Vara

## Abstract

Metabolic reprogramming is a hallmark of cancer. We and other authors have previously shown that breast cancer subtypes present metabolism differences. In this study, breast cancer cell lines were treated with metformin and rapamycin. The response was heterogeneous across various breast cancer cells, leading to cell cycle disruption in specific conditions. The molecular effects of these treatments were characterized using sublethal doses, SNP genotyping and mass spectrometry-based proteomics. Protein expression was analyzed using probabilistic graphical models, showing that treatments elicit various responses in some biological processes, providing insights into cell responses to metabolism drugs. Moreover, a flux balance analysis approach using protein expression values was applied, showing that predicted growth rates were comparable with cell viability measurements and suggesting an increase in reactive oxygen species response enzymes due to metformin treatment. In addition, a method to assess flux differences in whole pathways was proposed. Our results show that these various approaches provide complementary information, which can be used to suggest hypotheses about the drugs’ mechanisms of action and the response to drugs that target metabolism.

## Introduction

Reprogramming of cellular metabolism is a hallmark of cancer (Hanahan & Weinberg, 2011). Normal cells obtain energy mainly from mitochondrial metabolism, but cancer cells show increased glucose uptake and fermentation into lactate, which is known as the Warburg effect or aerobic glycolysis (Warburg, 1925). Cancer cells also exhibit increased glutamine uptake to maintain the pool of nonessential amino acids and to further increase lactate production (DeBerardinis et al, 2007).

Metabolic alterations enable the possibility of using metabolic inhibitors as targeted drugs. Metformin (MTF), a drug for diabetes, has begun clinical trials in cancer patients (Jones & Schulze, 2012). Everolimus, an inhibitor of mammalian target of rapamycin (mTOR), has clinical activity and has been approved for use in patients with breast cancer and other tumors (Beck, 2015). We previously observed significant differences in glucose metabolism between two of the main breast cancer subtypes: hormone-receptor positive (ER+) and triple-negative (TNBC) (Gámez-Pozo et al, 2015; Gámez-Pozo et al, 2017).

In the present study, we used single nucleotide polymorphism (SNP) profiling, proteomics and computational methods to explore the molecular consequences of metformin and rapamycin treatment in breast cancer cell lines. Additionally, protein expression data was included in a genome-scale model of metabolism and were analyzed using flux balance analysis (FBA).

High-throughput mass spectrometry-based proteomics allow the quantification of thousands of proteins and the acquisition of direct information about biological process effectors. Combined with probabilistic graphical models, proteomics enables the characterization of various biological processes between various conditions using expression data without other *a priori* information (Gámez-Pozo et al, 2015; Gámez-Pozo et al, 2017).

FBA is a widely used approach for modeling biochemical and metabolic networks in a genome scale (Edwards, 1999; Pramanik & Keasling, 1997; Varma & Palsson, 1995). FBA calculates the flow of metabolites through metabolic networks, allowing the prediction of growth rates or the rate of production of a metabolite. It has traditionally been used to estimate microorganism growth rates (Edwards et al, 2001). However, with the appearance of complete reconstructions of human metabolism, FBA has been applied to other areas such as red blood cells (Schilling & Palsson, 1998) or the study of the Warburg effect in cancer cell lines (Asgari et al, 2015).

Our results suggest that metformin and rapamycin treatments cause a heterogeneous effect on cell proliferation, with consequences at the cell cycle level. Functional analyses identified processes altered due to such treatments, and FBA predicted deregulation in reactive oxygen species (ROS) enzymes. We have shown that various analyses provide complementary information, which can be used to suggest hypotheses about the drugs’ mechanisms of action and responses that deserve subsequent validation.

## Results

### Breast cancer cell lines showed heterogeneous response when treated with drugs against metabolic targets

First, we evaluated the response of ER+ and TNBC breast cancer cell lines when treated with two drugs targeting metabolism, metformin (MTF) and rapamycin (RP). Cell viability was assessed for six breast cancer cell lines, three ER+ (T47D, MCF7 and CAMA1) and three TNBC (MDAMB231, MDAMB468 and HCC1143). Dose-response curves for each drug treatment in each cell were calculated (Table 1). A heterogeneous response was shown among breast cancer cell lines treated with a range of MTF and RP concentrations (Figure 1). Regarding RP, this heterogeneous response is related to breast cancer subtypes, showing an increased effect over ER+ cell line viability compared with those of TNBC.

**Figure 1:**
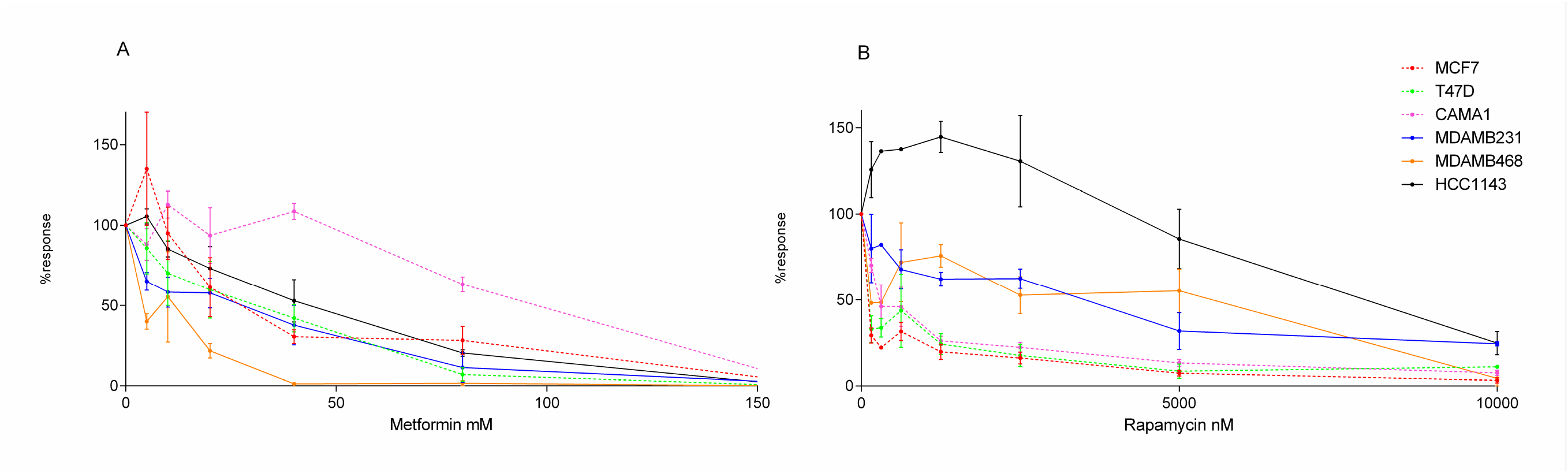
Dose-response curves of breast cancer cell lines treated with (A) MTF (0-160 mM) or (B) RP (0-10,000 nM). ER+ cell lines are represented as discontinuous lines and TNBC cells as continuous lines.

**Table 1:** Cell viability measurements in six breast cancer cell lines treated with MTF (0-160 mM) or RP (0-10,000 nM). Red-white-blue color scale.

| MTF mM | 0 | 5 | 10 | 20 | 40 | 80 | 160 |
| --- | --- | --- | --- | --- | --- | --- | --- |
| MCF7 | 100.00 | 135.07 | 95.00 | 61.49 | 30.61 | 28.36 | 2.47 |
| T47D | 100.00 | 85.74 | 70.15 | 59.87 | 42.11 | 7.10 | 0.00 |
| CAMA1 | 100.00 | 88.08 | 112.76 | 93.70 | 108.67 | 63.25 | 3.49 |
| MDAMB231 | 100.00 | 65.08 | 58.36 | 57.78 | 37.82 | 11.45 | 1.77 |
| MDAMB468 | 100.00 | 40.05 | 55.39 | 21.82 | 1.31 | 1.71 | 0.00 |
| HCC1143 | 100.00 | 105.48 | 85.25 | 73.19 | 52.89 | 20.49 | 0.00 |

| RP nM | 0 | 156.25 | 312.5 | 625 | 1250 | 2500 | 5000 | 10000 |
| --- | --- | --- | --- | --- | --- | --- | --- | --- |
| MCF7 | 100.00 | 29.36 | 22.34 | 31.62 | 19.88 | 16.29 | 7.53 | 3.32 |
| T47D | 100.00 | 33.02 | 33.76 | 43.74 | 24.39 | 17.73 | 8.69 | 11.15 |
| CAMA1 | 100.00 | 70.22 | 46.25 | 45.99 | 26.28 | 22.46 | 13.45 | 7.71 |
| MDAMB231 | 100.00 | 79.92 | 82.09 | 67.84 | 62.16 | 62.43 | 31.95 | 24.50 |
| MDAMB468 | 100.00 | 48.25 | 48.51 | 71.92 | 75.75 | 52.74 | 55.31 | 4.49 |
| HCC1143 | 100.00 | 125.74 | 136.39 | 137.53 | 144.66 | 130.58 | 85.55 | 24.85 |

### SNP genotyping of breast cancer cell lines

SNP genotyping was used to associate polymorphisms with the response of cell lines to MTF and RP. Polymorphisms previously related to drug sensitivity were studied using a custom expression array. Regarding the response to MTF, polymorphism rs2282143 in *SLC22A1* was detected in MDAMB468 cells. This SNP appears with a frequency of 8% in the black population, which is the population origin of this cell line, and it is associated with decreased clearance of MTF. On the other hand, the rs628031 polymorphism, also in *SLC22A1*, was found in homozygosis in MCF7 and HCC1143 cells and in heterozygosis with a possible duplication in MDAMB468 cells. The presence of this polymorphism was associated with a decreased response to MTF (PharmGKB; www.pharmgkb.org) (Sup Table 1).

Regarding the response to RP, two SNPs (rs1045642, rs2868177) in *ABCB1* and *POR* genes, respectively, were detected in hormone receptor-positive cell lines; rs1045642 was in heterozygosis in ER+ cell lines, and its effects are controversial. The relationship of rs2868177 with RP or another rapalog has not been previously described. MDAMB468 cells also present a polymorphism in heterozygosis in *CYP3A4* (rs2740574), which has been previously related to a requirement for an increased dose of RP as compared with a wild-type homozygote (PharmGKB; www.pharmgkb.org) (Sup Table 1).

### Molecular characterization of breast cancer cell lines’ response to treatment with drugs against metabolic targets using perturbation experiments and proteomics

When SNP genotyping did not fully explain the heterogeneous response between cell lines to MTF and RP treatment, we characterized the molecular basis of this heterogeneous response using proteomics in a perturbation experimental setting. Six breast cancer cell lines, treated or not with suboptimal concentrations of MTF and RP (40 mM of MTF [except for MDAMB468, in which a 20 mM concentration was used] and 625 nM of RP) were analyzed in duplicate using shotgun proteomics. Raw data normalization was performed, adjusting by duplicate values and as previously described (Gámez-Pozo et al, 2015). Mass spectrometry-based proteomics allowed the detection of 4052 proteins, which presented at least two unique peptides and detectable expression in at least 75% of the samples (Sup Table 2). No decoy protein passed through these additional filters. Label-free quantification values from these 4052 proteins were used in subsequent analyses.

We first identified proteins with differential expression between the treated and the control cells. Proteins with delta expression values between the control and treated cells higher than 1.5 or lower than -1.5 were identified for each cell line/treatment combination (Sup Tables 3 and 4). MCF7 cells treated with MTF showed decreased expression of 101 proteins, with a significant presence of proteins related to mitochondria and cell cycle, and increased expression of 52 proteins involved in mitochondria and cytoskeleton as majority functions. T47D cells treated with MTF presented decreased expression of 95 proteins and increased expression of 83 proteins, mostly related to mitochondria and the Golgi apparatus. CAMA1 treated with MTF had decreased expression of 105 proteins and increased expression of 53 proteins, without concrete functions overrepresented. MDAMB231 treated with MTF showed decreased expression of 135 proteins, mostly related to mitochondria, and an increase in 71 proteins. MDAMB468 presented decreased expression of 79 proteins, mostly related to mitochondria, and increased expression in 68 proteins mainly related to the extracellular matrix. Finally, HCC1143 showed decreased expression in 199 proteins, mostly related to mitochondria and mRNA processing, and increased expression in 58 proteins related to cytosol and protein binding.

Differentially expressed proteins were compared with gene interaction information contained in the Comparative Toxicogenomics Database. PIR, RELA, SIRT5, CMBL, PPP4R2 and MYD88 showed decreased expression, whereas SIRT2, SERPINE1 and HTATIP2 proteins showed increased expression in cells treated with MTF in both the database and in our experiments in at least one cell line.

Concerning RP treatment, MCF7 showed decreased expression in 98 proteins, mainly related to cellular transport, and an increased expression in 103 proteins mostly related to the mitochondrial matrix. T47D presented decreased expression in 115 proteins, most involved in cell division, and an increase in 82 proteins related to lysosomes. CAMA1 had a decrease in 452 proteins’ expression, mostly associated with mRNA processing, splicing and mitochondria, and an increase in 236 proteins, mostly associated with mitochondria, apoptosis processes and especially with the role of mitochondria in the apoptotic pathway. MDAMB231 had a decrease in 123 proteins related to mRNA processing and cytoskeleton, and an increase in 82 proteins related to exosomes. MDAMB468 had decreased expression in 58 proteins and increased expression in 82 proteins without a characteristic majority function. Lastly, HCC1143 showed a decreased expression in 103 proteins, mostly related to lysosomes, and an increased expression in 78 proteins, mostly related to mitochondria.

Gene interaction information contained in the Comparative Toxicogenomics Database showed a decrease in CDK4, CKS1B, COL1A1, IGFBP5, KIFC1, mTOR and SCD expression and an increase in CASP8, NR3C1, PKP4, RPS27L, TEAD1 and XIAP due to RP treatment in both the database and in our experiments in at least one cell line.

We applied linear regression models using protein expression data to discover molecular markers predicting the response to MTF and RP treatment. MMGT1, IDH1, PSPC1 and TACO1 showed the strongest correlation with the response to MTF (Sup Table 5); whereas ACADSB, CCD58, MPZL1 and SBSN correlated with the response to RP (Sup Table 6).

The next step was to explore molecular functions and biological pathways deregulated by MTF and RP treatment. Protein expression data from treated and untreated cells were used to build a probabilistic graphical model without other *a priori* information. The resulting graph was processed (Figure 2) to seek a functional structure, i.e., whether the proteins included in each branch of the tree had some relationship regarding their function, as previously described (Gámez-Pozo et al, 2015). Thus, we divided our graph into 36 branches and performed gene ontology analyses. Twenty-nine of them had a significant enrichment in proteins related to a specific biological function.

**Figure 2:**
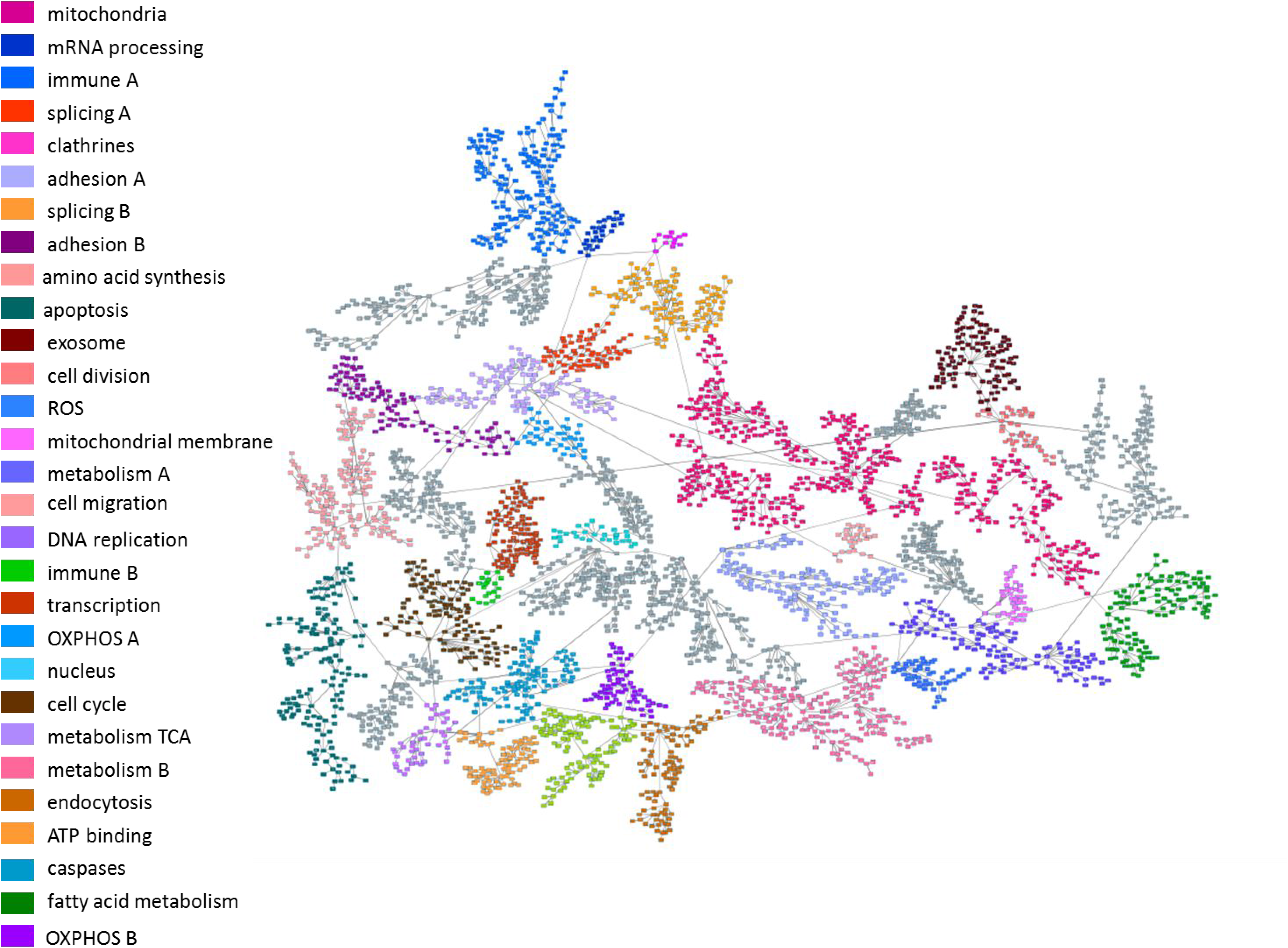
Probabilistic graphical model using protein expression data of control and treated breast cancer cell lines. Gray nodes lack a specific function.

Functional node activity was calculated for each branch with a defined biological function using protein delta values between control and treated cells. MTF treatment caused decreased activity in mitochondria B, mRNA processing, DNA replication and ATP binding functional nodes in all cell lines (Sup Fig 1). In the case of RP treatment, decreased activity was observed in mRNA processing node activity in all cell lines (Sup Fig 2).

Functional node activities were then evaluated using multiple linear regression models to explore the relationship between functional deregulation and MTF/RP treatment. The response to RP treatment was explained using metabolism A and B node activities (adjusted R^2^= 0.955). Metabolism A node is primarily related to fatty acid biosynthesis and pyrimidine metabolism and Metabolism B node is related to glycolysis, oxidative phosphorylation and carbon metabolism (Sup Table 7). The response to MTF could not be predicted using this approach.

### Cytometry experiments showed cytostatic effects of metformin and rapamycin treatment in breast cancer cells

The proteomics analysis workflow and gene ontology of delta values suggested that MTF and RP cause cell cycle alterations. To confirm this hypothesis, flow cytometry assessment of the cell cycle was performed. MCF7 and MDAMB231 cells treated with MTF showed an increased proportion of G2/M cells when compared with the control, suggesting a cell cycle arrest in the G2 phase. However, CAMA1 cells show an increase in G1 phase percentage. Regarding RP, the ER+ cell lines MCF7 and T47D treated with RP presented an increased percentage of G0/G1 cells when compared with the control, suggesting a cell cycle arrest in G1. On the other hand, the HCC1143 cycle showed an increase in G2 percentages (Figure 3, Sup Table 8).

**Figure 3:**
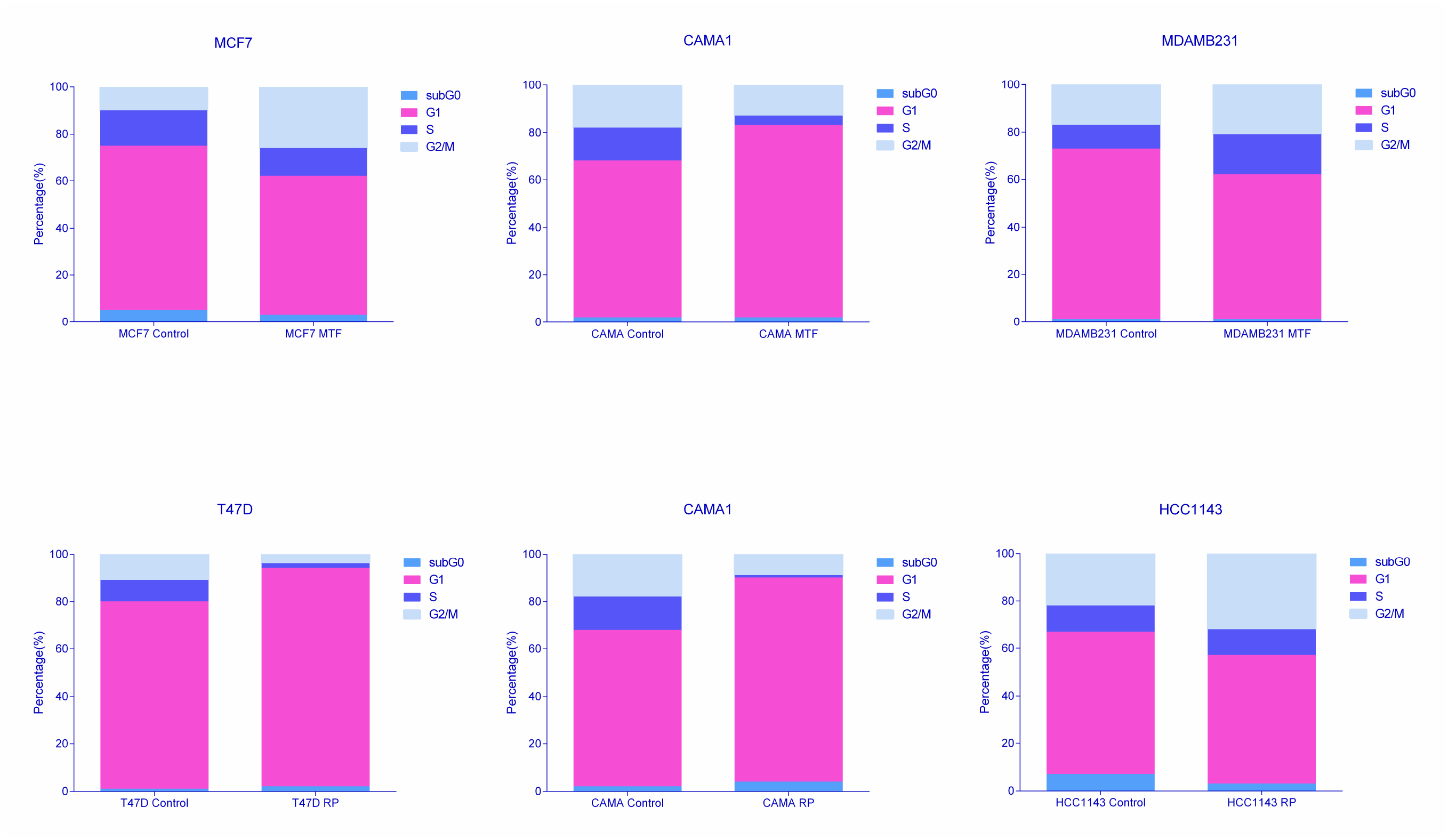
Percentages of cells in each cell cycle phase obtained by flow cytometry analyses.

### Flux balance analysis predicts alterations in growth rate in metformin-treated cells

FBA can be used to build a metabolic computational model that allows prediction of the tumor growth rate. This analysis can incorporate gene or protein expression data to improve prediction accuracy. To evaluate the impact of MTF and RP treatment on cellular metabolism, an FBA, including proteomics data from perturbation experiments, was applied to estimate cell growth rates for both control and treatment conditions. Protein data allows constraining 2414 reactions of the total number of 4253 reactions contained in Recon2, which has an associated gene-protein-reaction (GPR) rule. FBA predicts a lower growth rate in TNBC cells and MCF7 cell lines treated with MTF compared with control cells. However, it predicts a higher growth rate in the case of CAMA1 cells treated with MTF (Sup Table 9). FBA predicts no differences in growth rate between the control and the RP-treated cells.

### FBA growth predictions match with experimental data from breast cancer cell cultures

Growth kinetics studies in ER+ (MCF7 and T47D) and TNBC (MDAMB231 and MDAMB468) cell lines were performed to validate dynamic FBA cell growth model predictions. The starting concentration of glucose in medium was 200 mg/dl. This value was incorporated into the function inputs. Growth rate predictions were comparable with experimental measurements in cell cultures over 72 hours (Figure 4). The highest deviation in absolute values is observed in MDAMB468 cells, whereas MCF7 predictions coincided with experimental observations.

**Figure 4:**
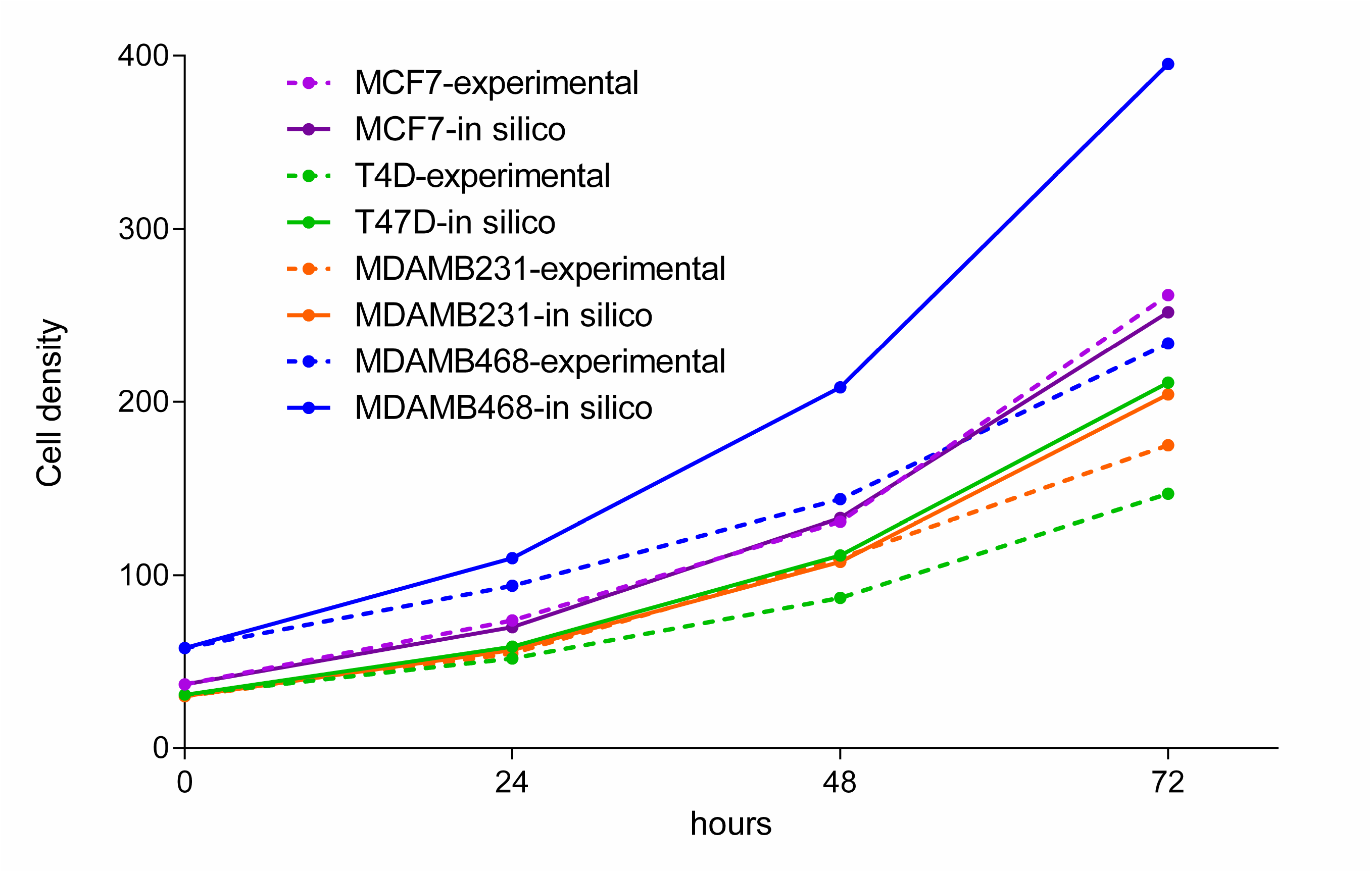
Experimental measurements of cell growth over 72 hours and a model simulation of growth during the same time period.

### Flux activity characterization

In order to compare pathway fluxes between untreated and treated cell lines, flux activities were calculated as the sum of the fluxes of each pathway. Pathways related to glutamate and pyruvate metabolism were related to response to MTF (adjusted R^2^=1) (Sup Table 10). In the case of RP, pathway fluxes that predict response against RP are cholesterol metabolism and valine, leucine and isoleucine metabolism (adjusted R^2^=1) (Sup Table 11).

### Flux analyses predict activation of ROS enzymes by metformin

With the aim of identifying reactions that changed as a consequence of treatment, we performed a Monte Carlo analysis and chose the solution with the maximum sum of fluxes because it was more representative of protein data (i.e., if a protein was measured, it indicated the protein was going to be used by the cell). After that, we used flux variability analysis (FVA) to calculate the possible maximum and minimum fluxes for each reaction, and therefore, the range of fluxes for each reaction. Next, we selected reactions showing a flux change between the control and the treated cells over 95% of this range. As long as FBA provides a unique optimal tumor growth rate, multiple combinations of fluxes can lead to this optimal value. Therefore, we confirmed that the results from the maximum flux solution were consistent throughout the multiple-solution landscape using a Monte Carlo approach to study a range of representative flux solutions from all possible solutions that optimize the tumor growth rate. Of all the candidates, we would like to highlight that FBA predicts a null catalase flux in control cells with the exception of HCC1143 cells, showing constitutive catalase activation. In MDAMB231 and MCF7 cell lines treated with MTF, the model predicts an activation of this reaction, whereas CAMA1 cells showed no response to MTF treatment regarding catalase activation (Sup Figure 3 and Sup Table 9).

Additionally, our model predicted that superoxide dismutase (SPODM) fluxes were increased in MCF7 and HCC1143 cell lines, but not in MDAMB231 cells. Predictions for CAMA1 cells showed high SPODM fluxes in both control and MTF treated cells (Sup Figure 4 and Sup Table 9).

Finally, the Monte Carlo approach predicted an increase in nitric oxide synthase flux and, as a consequence, an increase in nitric oxide (NO) production (Sup Figure 5).

On the other hand, proteomics data showed an increased expression of catalase in cells treated with MTF, with the exception of the CAMA1 cell line (Sup Table 8). Additionally, proteomics data showed an increased expression of SPODM in cells treated with MTF. However, in MDAMB231 cells, SPODM expression was generally lower than in the rest of the cell lines (Sup Table 12). No protein expression data from NO were obtained.

### Superoxide dismutase measurements confirm superoxide dismutase activation predictions

SPODM activities were measured in the control and in the MTF-treated cells using an enzyme activity assay. With the exception of the MCF7 cell line, model predictions were confirmed. In HCC1143, SPODM activity is similar between the control and the treated cells. On the other hand, MDAMB231 had the lowest SPODM activity, as shown in model predictions, and CAMA1 cells had the highest SPODM activity in the control and in the MTF-treated cells, as predicted in the model (Table 2).

**Table 2:** Superoxide dismutase activity assay measurements. The experiment was performed in triplicate, and one of the representative measurements is shown.

| Cell line | Superoxide dismutase activity (%) |
| --- | --- |
| MCF7 Control | 96.44% |
| MCF7 MTF | 90.76% |
| CAMA1 Control | 99.01% |
| CAMA1 MTF | 97.09% |
| MDAMB231 Control | 68.17% |
| MDAMB231 MTF | 49.82% |
| HCC1143 Control | 83.30% |
| HCC1143 MTF | 86.44% |

## Discussion

In this study, drugs targeting metabolism elicited changes related to cell cycle and oxidative stress in breast cancer cell lines. A high-throughput proteomics approach, coupled with a metabolism computational model, was useful to predict most of these changes and propose new mechanisms of action and effects of these drugs.

In previous studies, we observed significant differences between ER+ and TNBC glucose metabolism, which showed lactate production to be higher in TNBC cells than in ER+ cells (Gámez-Pozo et al, 2015). These metabolic alterations suggest the possibility of using drugs against metabolic targets in patients with breast cancer. SNP genotyping, proteomics, probabilistic graphical models and FBA in untreated and treated breast cancer cells were used to assess the mechanism of action and effects of metabolic drugs on breast cancer cells.

Our results show that breast cancer cells’ response to drugs targeting metabolism is heterogeneous. MTF treatment showed a broad effect on cell proliferation, with CAMA1 cells being the most resistant to this treatment. In the case of RP, the response depends on breast cancer subtype; it is effective in ER+ cell lines but not in those of TNBCs, resembling clinical results (a derivative of RP is used in women with hormone-receptor-positive breast cancer) (Beck, 2015).

With the aim of studying polymorphisms that could affect cell response, an SNP array was used. Therefore, sensitivity to MTF in MDAMB468 cells, which are the most affected by MTF, could be partly due to SNPs in the SLC22A1 carrier, which is related to decreased clearance of MTF. In addition, rs628031, previously associated with the poorest response against MTF, presented as homozygotic in the MCF7 and HCC1143 cell lines. ER+ cell lines presented heterozygosis in the ABCB1 rs1045642 polymorphism; however, the effects of this polymorphism are not yet clear. In CYP3A4, rs2740574, which is related to higher requirement of sirolimus, is shown as heterozygotic in the MDAMB468 cell line. Other polymorphisms were characterized, but none had been previously related to MTF or RP pharmacokinetics or pharmacodynamics.

In order to characterize the molecular changes provoked by MTF and RP treatments in breast cancer cell lines, a high-throughput proteomics approach was used to characterize perturbation experiments using these drugs. Next, differentially expressed proteins between the control and the treated cells were identified. Proteins related to response were also identified. Lastly, protein expression-based probabilistic graphical models were used to generate a functional structure, and differences in biological processes between the control and the treated cells were also characterized.

We discovered several differences between the MTF-treated cells and the control cells. Some of the differential proteins identified between the treated and the control cells matched with described interactions in the Comparative Toxicogenomics Database, such as increased expression of SIRT2 and HTATIP2 and decreased expression of SIRT5, PPP4R2 and MYD88 proteins due to MTF treatment. Increased SIRT2 protein expression induced by MTF treatment has been previously described (Buler et al, 2014). SIRT2 also enhances gluconeogenesis, plays an important inhibitory role in inflammation and elevates ROS defense (Gomes et al, 2015).

The effect of increased ROS stress response is in compliance with our model predictions. Moreover, MTF treatment results in decreased SIRT5 expression (Buler et al, 2014). This decrease is also related to differences observed in flux predictions between treated and control cells. It has been reported that SIRT5 is involved in the regulation of SPODM 1 activity (Lin et al, 2013), in accordance with our FBA prediction of SPODM activation in response to ROS stress in cells treated with MTF. On the other hand, TACO1, PSPC1, IDH1 and MMGT1 protein expression predict response to MTF treatment. IDH1 mutations were previously related to hypersensitivity to biguanides (Cuyàs et al, 2014). Probabilistic graphical models have shown that MTF treatment caused a decreased node activity in mRNA processing, DNA replication, mitochondria B and ATP binding nodes.

We also found several differences concerning RP treatment, such as an increased expression of NR3C1 and RPS27L proteins, and a decreased expression of CKS1B, COL1A1, IGFBP5, SCD, mTOR and CDK4 proteins, as previously reported (Tang et al, 2012). CDK4/6 inhibition robustly suppressed cell cycle progression of ER+/HER2-cellular models and complements the activity of limiting estrogen (Knudsen & Witkiewicz, 2016). RP treatment also results in decreased expression of CKS1B mRNA (Gonzalez et al, 2001). Knockdown of CKS1 expression promotes apoptosis of breast cancer cells (Wang et al, 2009). RP decreased expression of KIFC1 mRNA (Cui et al, 2011), whose overexpression is pro-proliferative (Pannu et al, 2015). RP treatment also results in increased activity of the NR3C1 protein (Davies et al, 2005). NR3C1 encodes the glucocorticoid receptor, which is involved in the inflammation response and which has an antiproliferative effect (Vilasco et al, 2011). RP enhances TP73 binding to the RPS27L promoter, a direct p53 target, and consequently promotes apoptosis (He & Sun, 2007). RP inhibits SCD mRNA expression through TP73 (Rosenbluth et al, 2011). 17-β-estradiol induces SCD expression and the modulation of cellular lipid composition in ER+ cell lines and is necessary for estrogen-induced cell proliferation (Belkaid et al, 2015). Finally, RP also decreases mTOR-related protein levels (Boulay et al, 2005; O'Reilly et al, 2011; Yee et al, 2006). Additionally, ACADSB, CCDC58, MPZL1 and SBSN protein expression predicts response to RP treatment. ACADSB affects valine and isoleucine metabolism (Andresen et al, 2000), which is one of the pathways related to response to RP in flux activity analyses, as we will explain later. Probabilistic graphical models showed that RP treatment caused decreased node activity in mRNA processing. Additionally, metabolism A and B node activities accurately predict the response in cells treated with RP.

Proteomics coupled with gene ontology analyses allowed us to explore protein expression between control and treated cells, suggesting that treatment with these drugs affects cell cycle progression. Therefore, the cell cycle was further assessed using flow cytometry. A cell cycle arrest in the G2/M phase was confirmed in all the MTF-treated cells except CAMA1, in which MTF had no effect on cell viability. Additionally, ER+ cells treated with RP (but not TNBC cells) had cell cycle arrest in G0/G1, which was confirmed at the cell proliferation level. It is known that mTOR controls cell cycle progression through S6K1 and 4E-BP1 (Fingar et al, 2004). Additionally, G0/G1 cell cycle arrest was previously described in MCF7 cells treated with RP (Tengku Din et al, 2014). Therefore, MTF and RP have cytostatic effects in breast cancer cell lines and cause a cell viability reduction, coupled with a disruption of the cell cycle. However, this response is diverse between various breast cancer cell lines.

On the other hand, FBA has been used in microbiology to study microorganism growth. This approach has recently been applied to study the Warburg effect (Asgari et al, 2015). We have developed a genome-scale cancer metabolic model that uses protein expression data to predict tumor growth rate. Previous studies have described cancer metabolic models using gene expression data (Asgari et al, 2015; Resendis-Antonio et al, 2010; Vázquez et al, 2010). Our model, however, used a whole human metabolism reconstruction and proteomics data to improve predictive accuracy. We assessed the model reliability by growth kinetics studies in ER+ (MCF7 and T47D) and TNBC (MDAMB231 and MDAMB468) cells. This approach allows new hypotheses and provides a global vision of metabolism, and has been previously used to characterize metabolism in samples from patients with breast cancer, which enables us to address clinically relevant questions (Gámez-Pozo et al, 2017).

Coupling proteomics data with FBA is challenging, primarily because relationships between protein expression data and enzymatic reactions is not direct. Data must be preprocessed through GPR rules to provide biologically meaningful information. On the other hand, functional interpretation of the data is not well established beyond individual changes in protein/reactions between control and treated scenarios. In this study, we overcame these limitations differently for proteins and reactions. Proteomics data were functionally explored using probabilistic graphical models. This tool has been used previously to identify functional differences using clinical samples (Gámez-Pozo et al, 2015; Gámez-Pozo et al, 2017). Regarding metabolic reactions, we propose the use of functional flux activities to compare complete pathways.

Model growth rate predictions were consistent with changes detected in viability assays in the cells treated with MTF. We explored the global flux for each pathway, calculating flux activities to identify metabolic pathways showing different behavior between the MTF-treated cells and the control cells. The pathways related to response to MTF treatment were glutamate and pyruvate metabolism. The pathways related to RP treatment response were valine, leucine and isoleucine metabolism, and cholesterol metabolism. Although it is difficult to make comparisons between flux patterns, pathway flux activities could be a useful approach to understanding changes between various conditions.

Moreover, by using an FVA coupled with the Monte Carlo approach, an activation of enzymes related to ROS stress response associated with MTF treatment could be predicted. Catalase and SPODM activation by MTF have been described in other scenarios (Dai et al, 2014; Kukidome et al, 2006), and as previously mentioned, concurs in most cases with differences shown in protein expression, although this relationship is not always direct. For instance, SPODM showed a 1.25-fold increase in protein expression, but no increment at the flux level, because fluxes are conditioned not only by their own restrictions, but also by bounds from adjacent reactions. In addition, catalase and SPODM fluxes appear to be related to cell viability. For instance, CAMA1 cells treated with MTF did not show an increased catalase flux, perhaps due to the discrete effect of MTF treatment on CAMA1 viability. Some of these predictions have been verified in the SPODM activity assay. In general, SPODM activity measurements were consistent with FBA predictions. Variations between FBA predictions and SPODM activities could be due to the fact that FBA only take into account metabolic pathways. On the other hand, our model predicts an increase in nitric oxide synthase flux in MCF7 cells treated with MTF, as has been previously described in diabetic rats (Volarevic et al, 2015). An increase in nitric oxide synthase implies a higher NO concentration, related to apoptosis processes and cytostatic effects in tumor cells, whereas low NO concentrations are associated with cell survival and proliferation (Vannini et al, 2015). This nitric oxide synthase activation could be related to the reduced proliferation observed in MCF7 cells treated with MTF. The fact that this effect was only predicted in MCF7 could be due to heterogeneity in the response mechanisms against this drug in various cellular contexts, and could be related to the observed differences in cell proliferation. It is remarkable that although no information about nitric oxide synthase abundance was provided by proteomics, our model reflects differences at the flux level in this process, suggesting that both approaches, proteomics and flux balance analysis, offer complementary information.

To summarize our results, treatment with MTF caused a heterogeneous effect on cell proliferation, consistent with a cell cycle arrest in the G2/M phase, and it appears to increase ROS enzymes. In MCF7 cells, an increase of nitric oxide synthase was predicted. Mitochondria and ATP binding node activities in the probabilistic graphical model are in compliance with the effect that MTF treatment has on mitochondria (El-Mir et al, 2000). Finally, susceptibility to MTF treatment shown by MDAMB468 cells could be related to an SLC22A1 SNP.

On the other hand, RP treatment exerts greater effect on the cell proliferation of ER+ cells, mediated by a G0/G1 cell cycle arrest, as previously described (Cuyàs et al, 2014). Our results suggest that valine and isoleucine metabolism could be deregulated by RP treatment. Finally, susceptibility of ER+ cell lines to RP treatment could be due to a SNP related to higher drug concentration.

Our study has some limitations. FBA provides an optimal biomass value, but multiple combinations of fluxes leading to this optimum are possible, making assessing differential pathways between conditions difficult. In our study, this limitation was solved using resampling techniques; however, improvement of computational processes is still necessary. Regarding proteomics experiments, although they can improve model accuracy, because they allow direct measurement of enzyme levels, at this moment this approach can only provide values for about 57% of Recon2 reactions with the known GPR rule. Gene expression, however, with the limitation of being an indirect measurement of enzyme abundance, provides almost the full picture. Strikingly, FBA was not able to reflect cell viability changes due to RP treatment. Despite the potential of the FBA approach, it only takes into account differences at the metabolic level. It is well known that mTOR inhibition leads to massive changes in cell homeostasis; thus, it appears reasonable that modeling changes at the metabolism level alone could not predict these differences.

We have characterized differential protein expression patterns between cells treated with drugs targeting metabolism and control cells. We have also developed a computational workflow to evaluate the impact of metabolic alterations in tumor and cell growth rates, using proteomics data. Growth rates predicted by our model matched the viability results observed *in vitro* with drug exposure. In addition, probabilistic graphical models are useful to study effects related to biological processes instead of considering individual protein or gene expression patterns. Our holistic approach shows that various analyses provide complementary information, which can be used to suggest hypotheses about drug mechanisms of action and response that deserve subsequent validation. Finally, this type of analysis, when fully developed and validated, could be used to study metabolic patterns from tumor samples with a different response against drugs targeting metabolism.

## Materials and Methods

### Cell culture and reagents

The ER+ breast cancer cell lines MCF7, T47D and CAMA1 and the triple-negative breast cancer cell lines MDAMB231, MDAMB468 and HCC1143 were cultured in RPMI-1640 medium with phenol red, supplemented with 10% heat-inactivated fetal bovine serum, 100 mg/mL penicillin and 100 mg/mL streptomycin. All the cell lines were cultured at 37°C in a humidified atmosphere with 5% (v/v) CO_2_ in the air. The MCF7, T47D and MDA-MB-231 cell lines were kindly provided by Dr. Nuria Vilaboa (La Paz University Hospital, previously obtained from ATCC in January 2014). The MDAMB468, CAMA1 and HCC1143 cell lines were obtained from ATCC (July 2014). Cell lines were routinely monitored in our laboratory and authenticated by morphology and growth characteristics, tested for Mycoplasma and frozen, and passaged for fewer than 6 months before experiments. The MTF (Sigma Aldrich D150959) and RP (Sigma Aldrich R8781) were obtained from Sigma-Aldrich (St. Louis, MO, USA).

### Cell viability assays

The cells were treated with MTF and RP at a range of concentrations to establish an IC_50_ for each cell line. Approximately 5000 cells per well were seeded in 96-well plates. After 24 h, an appropriate concentration of drug was added to the cells, which were incubated for a total of 72 h. Untreated cells were used as a control. The CellTiter 96 AQueous One Solution Cell Proliferation Assay (Promega) kit was used for the quantification of cell survival after exposure to the drugs. After 72 h of incubation with the drug, CellTiter 96 AQueous One Solution was added to each well following the manufacturer’s instructions, and absorbance was measured on a microplate reader (TECAN). Experiments were performed in triplicate. IC_50_ values were calculated using the Chou-Talalay method (Chou, 2006).

### SNP genotyping

We used TaqMan OpenArray technology on a QuantStudio 12K Flex Real-Time PCR System (Applied Biosystems^®^) with a custom SNP array format, which allows simultaneous genotyping of 180 SNPs in major drug metabolizing enzymes and transporters (PharmArray^®^). Information about the pharmacogenetic variants associated with RP and MTF response was gathered mostly from the variant and clinical annotations in the Pharmacogenomics Knowledge Base (PharmGKB; www.pharmgkb.org). The final selection of SNPs for our study was as follows: rs2032582, rs1045642, rs3213619 and rs1128503 in the *ABCB1* gene; rs55785340, rs4646438 and rs2740574 in *CYP3A4;* rs776746, rs55965422, rs10264272, rs41303343 and rs41279854 in *CYP3A5;* rs1057868 and rs2868177 in *POR* for RP; and rs55918055, rs36103319, rs34059508, rs628031, rs4646277, rs2282143, rs4646278, rs12208357 in *SLC22A1* and rs316019, rs8177516, rs8177517, rs8177507 and rs8177504 in *SLC22A2* for MTF. Molecular analyses for rs34130495 and rs2740574 were performed by classic sequencing because these probes were not originally included in our custom SNP array design.

### Perturbation experiments

Suboptimal concentrations (IC_70_ or higher) were chosen in order to perform perturbation experiments (MTF 40 mM except for MDAMB468 20 m, RP 625 nM). Approximately 500,000 cells per well were seeded in 6-well plates. Twenty-four hours later, drugs against metabolism were added. After additional 24 h, proteins were extracted using the ISOLATE II RNA/DNA/Protein Kit (BIOLINE). Protein concentration was determined using the MicroBCA Protein Assay Kit (Pierce-Thermo Scientific). Protein extracts (10 μg) were digested with trypsin (Promega) (1:50). Peptides were desalted using in-house-produced C18 stage tips, then dried and resolubilized in 15 μl of 3% acetonitrile and 0.1% formic acid for MS analysis.

### Liquid chromatography - mass spectrometry shotgun analysis

Mass spectrometry analysis was performed on a Q Exactive mass spectrometer coupled to a nano EasyLC 1000 (Thermo Fisher Scientific). Solvent composition at the two channels was 0.1% formic acid for channel A; and 0.1% formic acid, 99.9% acetonitrile for channel B. For each sample, 3 μL of peptides were loaded on a self-made column (75 μm × 150 mm) packed with reverse-phase C18 material (ReproSil-Pur 120 C18-AQ, 1.9 μm, Dr. Maisch GmbH) and eluted at a flow rate of 300 nL/min at a gradient from 2% to 35% B in 80 min, 47% B in 4 min and 98% B in 4 min. Samples were acquired in a randomized order. The mass spectrometer was operated in data-dependent mode, acquiring a full-scan MS spectra (300–1700 m/z) at a resolution of 70,000 at 200 m/z after accumulation to a target value of 3,000,000, followed by higher-energy collisional dissociation (HCD) fragmentation on the 12 most intense signals per cycle. The HCD spectra were acquired at a resolution of 35,000 using normalized collision energy of 25 and a maximum injection time of 120 ms. The automatic gain control was set to 50,000 ions. Charge state screening was enabled, and single and unassigned charge states were rejected. Only precursors with intensity above 8300 were selected for MS/MS (2% underfill ratio). Precursor masses previously selected for MS/MS measurement were excluded from further selection for 30 s, and the exclusion window was set at 10 ppm. The samples were acquired using internal lock mass calibration on m/z 371.1010 and 445.1200.

### Protein identification and label-free protein quantification

The acquired raw MS data were processed by MaxQuant (version 1.4.1.2), followed by protein identification using the integrated Andromeda search engine. Each file is kept separate in the experimental design to obtain individual quantitative values. The spectra were searched against a forward Swiss-Prot human database, concatenated to a reversed decoyed FASTA database and common protein contaminants (NCBI taxonomy ID9606, release date 2014-05-06). Methionine oxidation and N-terminal protein acetylation were set as variable modification. Enzyme specificity was set to trypsin/P allowing a minimal peptide length of 7 amino acids and a maximum of two missed cleavages. Precursor and fragment tolerance was set to 10 ppm and 20 ppm, respectively, for the initial search. The maximum false discovery rate (FDR) was set to 0.01 for peptides and 0.05 for proteins. Label-free quantification was enabled, and a 2-minute window for match between runs was applied. The requantify option was selected. For protein abundance, the intensity (Intensity) as expressed in the protein groups file was used, corresponding to the sum of the precursor intensities of all identified peptides for the respective protein group. Only quantifiable proteins (defined as protein groups showing two or more razor peptides) were considered for subsequent analyses. Protein expression data were transformed (hyperbolic arcsine transformation), and missing values (zeros) were imputed using the missForest R package (Stekhoven & Bühlmann, 2012). The protein intensities were normalized by scaling the median protein intensity in each sample to the same values. Then values were log_2_ transformed.

### Gene ontology analyses

Protein expression patterns were compared between the control and treated cells, and deltas were calculated for each drug in each cell line by subtracting control protein expression from treated cell protein expression values. Gene ontology analyses were performed to determine differential functions between the control and the treated cells. For this, we selected protein showing a change in expression values (delta) higher than 1.5 or lower than -1.5; this delta value was calculated for each protein as the treated cell expression value minus the control cell expression value. Protein-to-gene ID conversion were performed using Uniprot (http://www.uniprot.org) and DAVID (Huang et al, 2009). The gene ontology analyses were performed using the functional annotation chart tool provided by DAVID. We used “homo sapiens” as a background list and selected only GOTERM-FAT gene ontology categories and Biocarta and KEGG pathways. Functional categories with p<.05 and a FDR below 5% were considered as significant.

### Probabilistic graphical models, functional node activity measurements and response predicted models

Network construction was performed using probabilistic graphical models compatible with high dimensional data using correlation coefficients as associative measures as previously described (Gámez-Pozo et al, 2015).

The network was split into several branches, and a gene ontology analysis was used to explore the major biological function for each branch, defining functional nodes. Functional node activity was calculated as the mean delta, between treated and untreated cells, of all proteins related to the assigned majority node function. In order to relate drug response to functional processes, multiple linear regression models were performed using IBM SPSS Statistics.

### Cytometry experiments

Some 500,000 cells were seeded in each well per duplicate. Twenty-four hours later, drugs were added, and after 72 h, the cells were fixed in ethanol and marked with propidium iodide. Cells were acquired using a FACScan cytometer equipped with a blue laser at a wavelength of 488 nm. Acquired data were analyzed using BD CellQuest Pro software, first filtering cells by size and complexity in order to exclude debris, and then excluding doublets and triplets by FL2-W/FL2-A.

### Flux balance analysis and E-flux algorithm

FBA was used to build a metabolic computational model that predicts growth rates. FBA calculates the flow of metabolites through metabolic networks and predicts growth rates or the rate of production of a given metabolite. It was performed using the COBRA Toolbox (Schellenberger et al, 2011) available for MATLAB and the human metabolism reconstruction Recon2 (Thiele et al, 2013). The biomass reaction proposed in Recon2 was used as an objective function representative of growth rate in tumor cells. Proteomics expression data were included in the model by solving GPR rules and the E-flux algorithm (Colijn et al, 2009). Measuring GPR rule estimation values was performed using a variation of the method described by Barker et al (Barker et al, 2015). The mathematical operations used to calculate the numerical value were the sum of “OR” expressions and the minimum of “AND” expressions. Finally, the GPR rule values, *aj*, were normalized to a [0, 1] interval, using a uniform distribution formula. The normalized values have been used to establish both new lower and upper reaction bounds. If the reaction is irreversible the new bounds are 0 and *aj*, and if the reaction is reversible the new bounds are - *aj* and *aj.*

### Metabolism model validation

In order to validate model predictions we used dynamic FBA, which allows the prediction of cell growth during a period of time (Resendis-Antonio et al, 2010), and kinetic studies of cell lines were performed. This consists of an iterative approach based on a quasi-steady state assumption (Varma & Palsson, 1994). MCF7, T47D, MDAMB468 and MDAMB231 were seeded at an initial cell density of 1,000,000 cells. Cells within the same area were counted once a day for 3 days. To perform the dynamic FBA, experimental cell density at the beginning and experimental measured glucose concentration in the medium were used as inputs in the computational simulation. Glucose presented in the medium was measured using an ABL90 FLEX blood analyzer (Radiometer).

### Flux activities

With the aim of comparing the activity of the various pathway fluxes between the control and the treated cells, flux activity was calculated for each condition. Flux activity was defined by the sum of fluxes for all reactions involved in one pathway as defined in the Recon2. Then, linear regression models were performed.

### Flux variability analysis and the Monte Carlo approach

One obvious limitation to the FBA approach is that this analysis provides a unique optimal tumor growth rate, however, multiple combinations of fluxes can lead to this optimal value. In order to evaluate a representative sample of these multiple solutions, a Monte Carlo approach (Schellenberger, 2010) was used to compare differential fluxes between treated and untreated cells. The solution showing the maximum sum of all the fluxes was then used to calculate the flux change between the control and the treated cells. This criterion was selected under the premise that if a protein was experimentally measured it was because that protein was going to be used by the cell; thus, maximum flux solution picks up all measured proteins. On the other hand, FVA provides the possible maximum and minimum fluxes for each reaction; therefore, the flux range for each reaction. This range was used to calculate the flux change between the control and the treated cells for a given reaction as a percentage of the flux range for that reaction. Reactions showing a flux change between the control and the treated cells over 95% of this range were identified for each condition. Monte Carlo results for these reactions were used to check if maximum solution flux is representative of the most frequent solution flux for this reaction.

### Superoxide dismutase activity assay

To validate some of our model hypotheses, a SPODM activity assay was performed in triplicate, using the Superoxide Dismutase Assay Kit (Sigma-Aldrich, 19160). Some 500,000 cells per well were seeded, and after 24 h, MTF was added at 40 mM (except for the MDAMB468 cell line, in which a 20 mM concentration was used). Twenty-four hours later, SPODM activities were measured following the manufacturer’s instructions.

### Statistical analyses and software suites

Dose-response curves were constructed with GraphPad Prism 6. Gene and protein interactions for each drug were obtained from the Comparative Toxicogenomics Database (http://ctdbase.org/") (Davis et al, 2017). Linear and multiple regression models were built using IBM SPSS Statistics.

## Acknowledgments

This study was supported by Instituto de Salud Carlos III, Spanish Economy and Competitiveness Ministry, Spain and co-funded by the FEDER program, “Una forma de hacer Europa” (PI15/01310). LT-F is supported by the Spanish Economy and Competitiveness Ministry (DI-15-07614). The funders had no role in the study design, data collection and analysis, decision to publish or preparation of the manuscript. The cytometry experiments were performed at the Cytometry and Fluorescence Microscopy Center, Faculty of Chemistry, Complutense University of Madrid, Spain.

## Author contributions

All the authors have directly participated in the preparation of this manuscript and have approved the final version submitted. LT-F and RL-V performed the viability experiments and prepared the proteomics samples. LT-F performed the FBA. PN contributed the mass spectrometry data. JMA, HN and PM contributed the probabilistic graphical models. MD-A contributed the GPR rule calculation. LT-F, RL-V and RA-L contributed the enzyme activity assays. LT-F, GP-V, AZ-M and SL-A performed the statistical analysis, the probabilistic graphical model interpretation and the gene ontology analyses. ID, PA and AMB contributed the SNP study. LT-F drafted the manuscript. LT-F, AG-P, JAFV, JF and EE conceived of the study and participated in its design and interpretation. AG-P, JAFV, and EE supported the manuscript drafting. AG-P and JAFV coordinated the study. All the authors have read and approved the final manuscript.

